# MAST: A flexible statistical framework for assessing transcriptional changes and characterizing heterogeneity in single-cell RNA-seq data

**DOI:** 10.1101/020842

**Authors:** Greg Finak, Andrew McDavid, Masanao Yajima, Jingyuan Deng, Vivian Gersuk, Alex K. Shalek, Chloe K. Slichter, Hannah W. Miller, M. Julianna McElrath, Martin Prlic, Peter S. Linsley, Raphael Gottardo

**Affiliations:** Vaccine and Infectious Disease Division, Fred Hutchinson Cancer Research Center, Seattle, WA 98109, USA; Public Health Sciences Division, Fred Hutchinson Cancer Research Center and Benaroya Research Institute at Virginia Mason, Seattle, WA 98101, USA; Institute for Medical Engineering & Science & Department of Chemistry, MIT, Boston, MA, 01239-4307, USA; Ragon Institute of MGH, MIT, & Harvard, Boston, MA, 02139-3583, USA; Broad Institute of MIT & Harvard, Boston, MA, 01242, USA

## Abstract

Single-cell transcriptomic profiling enables the unprecedented interrogation of gene expression heterogeneity in rare cell populations that would otherwise be obscured in bulk RNA sequencing experiments. The stochastic nature of transcription is revealed in the bimodality of single-cell transcriptomic data, a feature shared across single-cell expression platforms. There is, however, a paucity of computational tools that take advantage of this unique characteristic. We present a new methodology to analyze single-cell transcriptomic data that models this bimodality within a coherent generalized linear modeling framework. We propose a two-part, generalized linear model that allows one to characterize biological changes in the proportions of cells that are expressing each gene, and in the positive mean expression level of that gene. We introduce the *cellular detection rate*, the fraction of genes turned on in a cell, and show how it can be used to simultaneously adjust for technical variation and so-called “extrinsic noise” at the single-cell level without the use of control genes. Our model permits direct inference on statistics formed by collections of genes, facilitating gene set enrichment analysis. The residuals defined by such models can be manipulated to interrogate cellular heterogeneity and gene-gene correlation across cells and conditions, providing insights into the temporal evolution of networks of co-expressed genes at the single-cell level. Using two single-cell RNA-seq datasets, including newly generated data from Mucosal Associated Invariant T (MAIT) cells, we show how model residuals can be used to identify significant changes across biologically relevant gene sets that are missed by other methods and characterize cellular heterogeneity in response to stimulation.

## Introduction

Whole transcriptome expression profiling of single cells via RNA-seq (scRNA-seq) is the logical apex to single cell gene expression experiments. In contrast to transcriptomic experiments on mRNA derived from bulk samples, this technology provides powerful multi-parametric measurements of gene co-expression at the single-cell level. However, the development of equally potent analytic tools has trailed the rapid advances in the biochemistry and molecular biology, and several challenges need to be addressed to fully leverage the information in single-cell expression profiles.

First, single-cell expression has repeatedly been shown to exhibit a characteristic bimodal expression pattern, wherein the expression of otherwise abundant genes is either strongly positive, or undetected within individual cells. This is due in part to low starting quantities of RNA such that many genes will be below the threshold of detection, but there is also a biological component to this variation (termed extrinsic noise in the literature) that is conflated with the technical variability^1-3^. We and other groups^4-6^ have shown that the proportion of cells with detectable expression reflects both technical and biological differences between samples. Results from synthetic biology also support the notion that bimodality can arise from the stochastic nature of gene expression^2,3,7,8^.

Secondly, measuring single cell gene expression might seem to obviate the need to normalize for starting RNA quantities. Recent work shows that cells scale transcript copy number with cell volume (a factor that affects gene expression globally) to maintain a constant mRNA concentration and thus constant biochemical reaction rates^9,10^. In scRNA-seq, cells of varying volume are diluted to an approximately fixed reaction volume leading to differences in detection rates of various mRNA species that are driven by the initial cell volumes. Technical assay variability (e.g. mRNA quality, pre-amplification efficiency) and extrinsic biological factors (e.g. nuisance biological variability due to cell size) remain, and can significantly influence expression level measurements. Consequently, this may render traditional normalization strategies using the expression level of a few “housekeeping” genes, like GAPDH, infeasible^10^. Recently, Shalek et al^5^ observed a strong relationship between average expression and detection efficiency, and have proposed a computational approach to correct the estimated gene-specific probability of detection. Our approach easily allows for estimation and control of the CDR simultaneously while estimating treatment effects as opposed to previous approaches^5^ that relied on a set of control genes and could not jointly model both factors.

Previously, Kharchenko et al^6^ developed a so-called three-component mixture model to test for differential gene expression while accounting for bimodal expression. Their approach is limited to two-class comparisons and cannot adjust for important biological covariates such as multiple treatment groups and technical factors such as batch or time information, severely limiting its utility in more complex experimental designs. On the other hand, several methods have been proposed for modeling bulk RNA-seq data that permit complex modeling through linear^11^ or generalized linear models^12,13^ but these models have not yet been adapted to single-cell data as they do not properly account for the observed bimodality in expression levels. This is particularly important when adjusting for covariates that might affect the expression rates. As we will demonstrate later, such model mis-specification can significantly affect sensitivity and specificity when detecting differentially expressed genes and gene-sets.

Here, we propose a Hurdle model tailored to the analysis of scRNA-seq data, providing a mechanism to address the challenges noted above. It is a two-part generalized linear model that simultaneously models the rate of expression over background of various transcripts, and the positive expression mean. Leveraging the established theory for generalized linear modeling allows us to accommodate complex experimental designs while controlling for covariates (including technical factors) in both the discrete and continuous parts of the model. We introduce the *cellular detection rate (CDR)*: the fraction of genes that are turned on / detected in each cell, which, as discussed above, acts as a proxy for both technical (e.g. dropout, amplification efficiency, etc.) and biological factors (e.g. cell volume and other extrinsic factors other than treatment of interest) that can influence gene expression. As a result it represents an important source of variability in scRNA-seq data that needs to be considered (Figure 1). Our approach of modeling the CDR as a covariate, offers an alternative to the weight correction of Shalek et al^5^ that does not depend on the use of control genes and allows us to jointly estimate nuisance and treatment effects. Our framework permits the analysis of complex experiments, such as repeated single cell measurements under various treatments and/or longitudinal sampling of single cells from multiple subjects with a variety of background characteristics (e.g. gender, age, etc.) as it is easily extended to accommodate random effects. Differences between treatment groups are summarized with pairs of regression coefficients whose sampling distributions are available through bootstrap or asymptotic expressions, enabling us to perform complementary differential gene expression and gene set enrichment analyses (GSEA). We use an empirical Bayesian framework to regularize model parameters, which helps improve inference for genes with sparse expression, much like what has been done for bulk gene expression^14^. Our GSEA approach accounts for gene-gene correlations, which is important for proper control of type I errors^15^. This GSEA framework is particularly useful for synthesizing observed gene-level differences into statements about pathways or modules. Finally, our model yields *single cell residuals* that can be manipulated to interrogate cellular heterogeneity and gene-gene correlations across cells and conditions. We have named our approach MAST for Model-based Analysis of Single-cell Transcriptomics.

**Figure 1.**
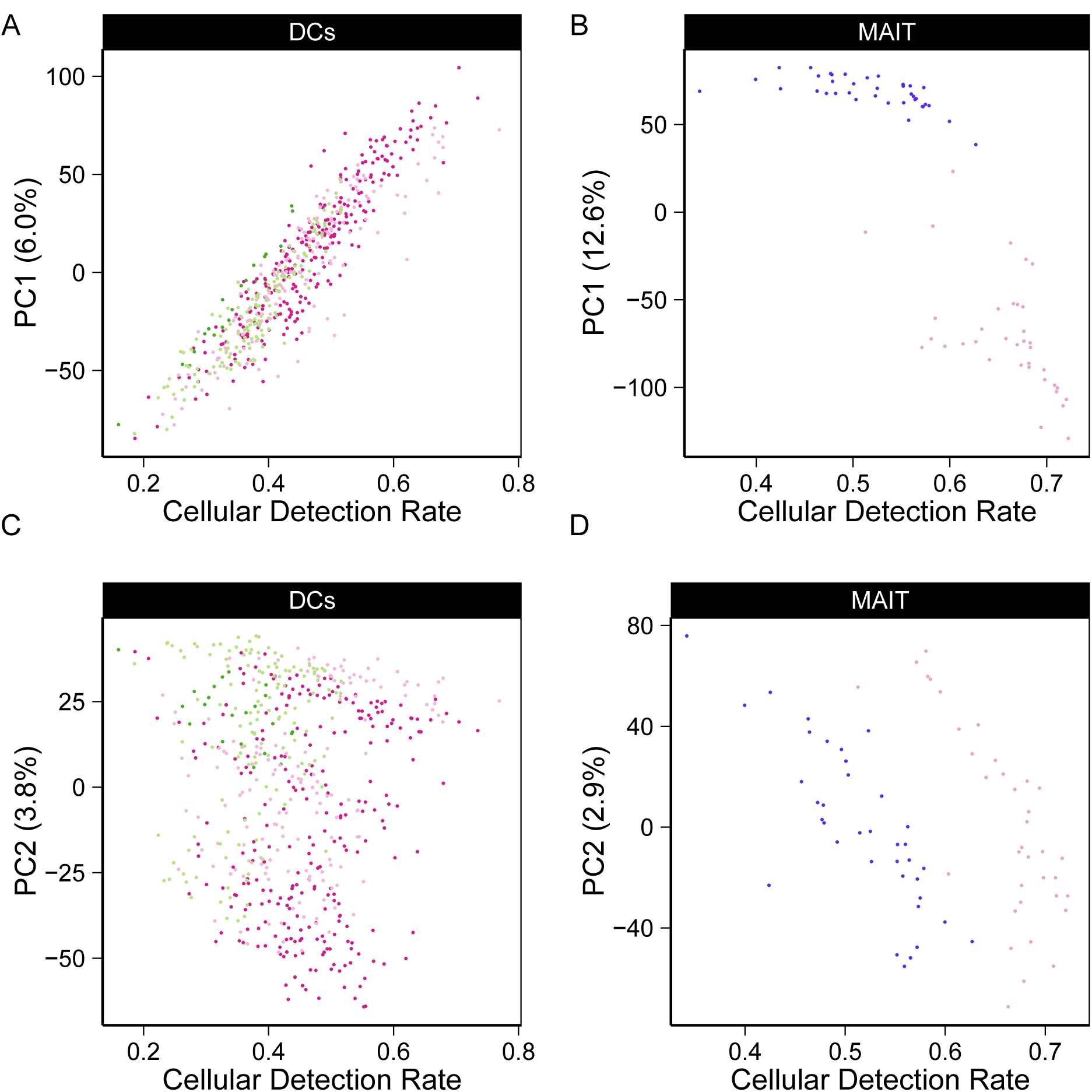
The fraction of genes expressed, or cellular detection rate (CDR), explains the principal components of variation in MAIT and DC data sets.

We illustrate the method on two data sets. We first apply our approach to an experiment comparing primary human non-stimulated and cytokine-activated Mucosal-Associated Invariant T (MAIT) cells. MAST identifies novel expression signatures of activation, and the single-cell residuals produced by the model highlights a population of MAIT cells showing partial activation but no induction of effector function. We then illustrate the application of MAST to a previously-published complex experiment studying temporal changes in murine bone marrow-derived dendritic cells subjected to LPS stimulation. We both recapitulate the findings of the original publication and describe additional coordinated gene expression changes at the single-cell level across time in LPS stimulated mDC cells.

## Results

### MAST can account for variation in the cellular detection rate

As discussed previously and as shown on Figure 1 by principal component analysis (PCA), the cellular detection rate (CDR, see Methods for exact definition), is an important source of variability. It is highly correlated with the second principal component (PC, Pearson’s rho=0.76 grouped, 0.91 stimulated, 0.97 non-stimulated) in the MAIT dataset and the first PC (rho=0.92 grouped, 0.97 non-stimulated, 0.92 LPS, 0.89 PAM, 0.92 PIC) in the mDC dataset. We observe larger CDR variability within treatment groups than across groups, suggesting that it is likely to be a nuisance factor. This is further supported by the fact that the CDR calculated within control (e.g. housekeeping) genes is highly correlated with the CDR calculated over all genes (Supplementary Figure 1). Its role as a principal source of variation persists across experiments (Figure 1).

We thus conjecture that CDR is a proxy for unobserved nuisance factors that should be explicitly modeled. In particular, it is not unreasonable to suggest that the CDR captures variation in global transcription rate due to variations in cell size (among other factors)^10^, as well as technical variation such as dropout, with dropout rates possibly correlated with cell-size. Fortunately, MAST easily accommodates covariates, such as the CDR, and more importantly allows joint, additive modeling of them with other biological variables of interest, with the effect of each covariate decomposed into its discrete and continuous parts. This two-part modeling is key to account for the CDR that directly reflects the gene-level transcription rates. Applying an analysis of deviance with MAST (see Methods), we quantified the amount of variability that could be attributed to CDR. The CDR accounts for 5.2% of the deviance in the MAIT data set and 4.8% in the mDC data set for the average gene, and often times much more than that: it comprises more than 9% of the deviance in over 10% of genes in both data sets, particularly for the discrete component of the model (Supplementary Figure 2). It should also be noted that the CDR deviance estimates for many of the genes are comparable (if not greater) to the treatment deviance estimates showing that it.

**Figure 2.**
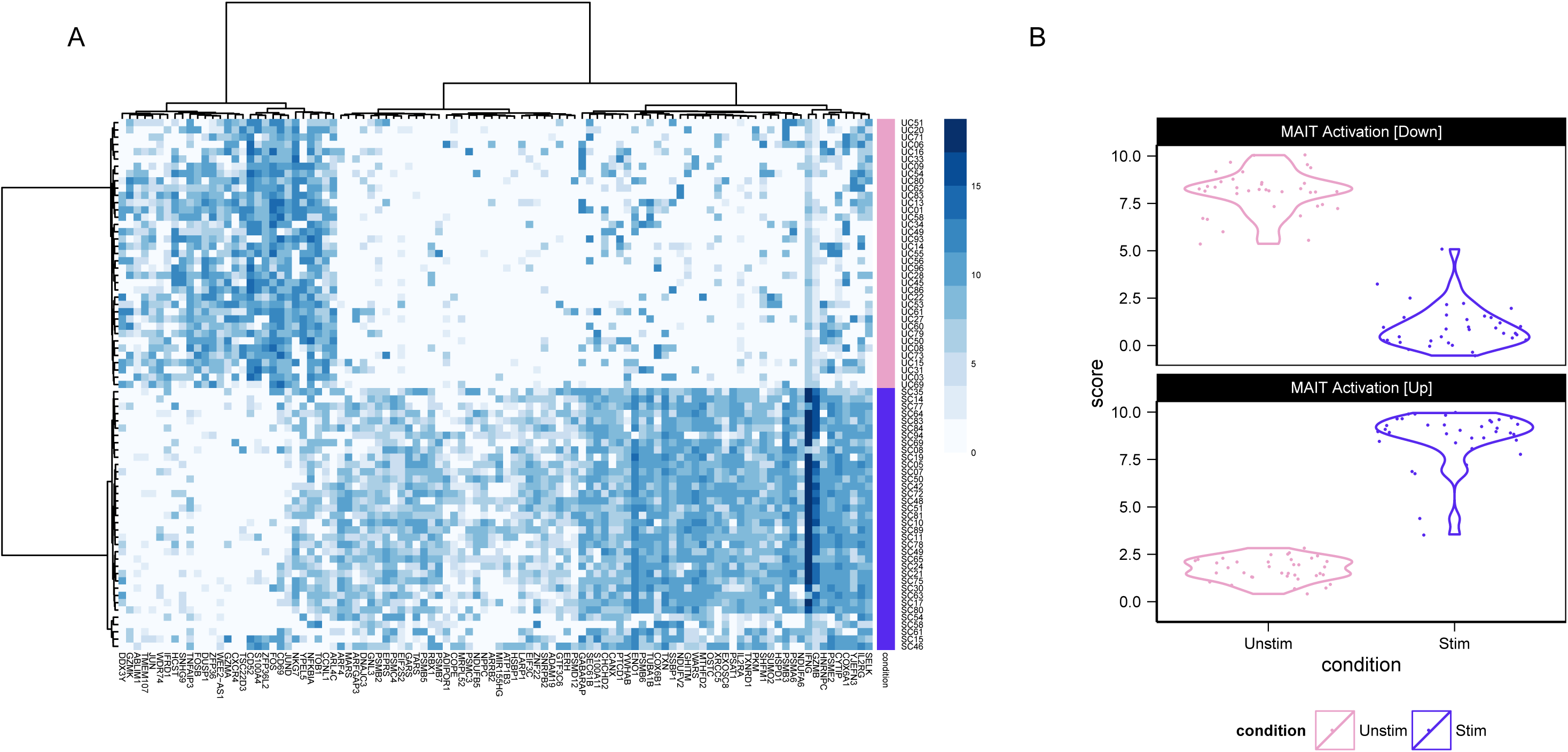
Single-cell expression (log_2_-TPM) of the top 100 genes identified as differentially expressed between cytokine (IL18, IL15, IL12) stimulated (purple) and non-stimulated (pink) MAIT cells using MAST (A). Partial residuals for up- and down-regulated genes are accumulated to yield an activation score (B), and this score suggests that the stimulated cells have a more heterogeneous response to stimulation than do the non-stimulated cells.

That CDR predicts expression levels contradicts the model of independent expression between genes, since the level of expression (averaged across many genes) would not affect the level in any given gene were expression independent. This pervasiveness suggests latent factors are creating coordinated changes in expression across genes. In light of the work of Padovan-Merhar et al^10^, we conjecture the latent factor relates to differences in cell volumes, since cells of different volumes compensate to conserve mRNA species molarity, which implies higher copy numbers of all transcripts in larger cells. Higher copy numbers result in higher scRNA-seq detection rates globally across transcripts.

Finally, we have investigated the relationship between our approach and the weight correction of Shalek et al^5^ (Supplementary Figure 3). We observe a strong linear relationship between the CDR and the weights of Shalek et al^5^. Thus, use of the CDR as a covariate can be seen as a statistically rigorous way to correct for the dropout biases of Shalek et al^5^, without the need to use control genes, and more importantly with the ability to control for these while estimating treatment effects.

**Figure 3.**
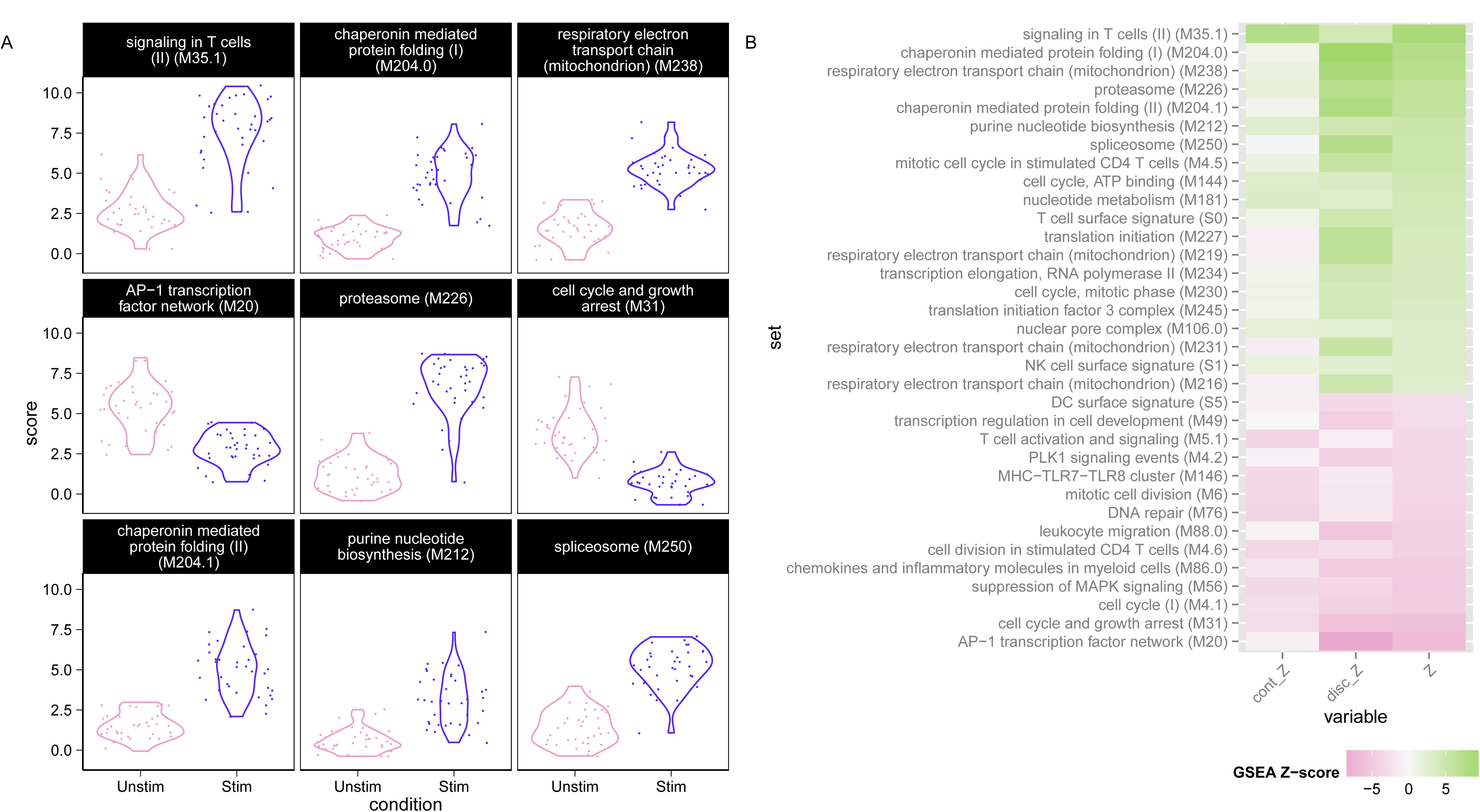
Module scores for individual cells for the top 9 enriched modules (A) and decomposed Z-scores (B) for single-cell gene set enrichment analysis in MAIT data set, using the blood transcription modules (BTM) database. The distribution of module scores suggests heterogeneity among individual cells with respect to different biological processes. Enrichment of modules in stimulated and non-stimulated cells is due to a combination of differences in the discrete (proportion) and continuous (mean conditional expression) of genes in modules. The combined Z-score reflects the enrichment due to differences in the continuous and discrete components.

### Single-cell sequencing identifies a transcriptional profile of MAIT cell activation

We applied MAST to our MAIT dataset to identify genes up- or down-regulated by cytokine stimulation while accounting for variation in the CDR (see Methods). We detected 291 differentially expressed genes, as opposed to 1413 when excluding CDR. To determine whether this was due to a change in ranking or a simply a shift in significance, we compared the overlap between the top *n* genes in both models (varying *n* from 100 to 1413), and found that, on average, 35% (range 32% - 38%) of genes are excluded when CDR is modeled, suggesting that inclusion of this variable allows global changes in expression, manifest in the CDR, to be decomposed from local changes in expression. This is supported by gene ontology enrichment analysis (Supplementary Figure 4) of these CDR-specific genes (n=539), where we see no enrichment for modules associated with treatment of interest.

**Figure 4.**
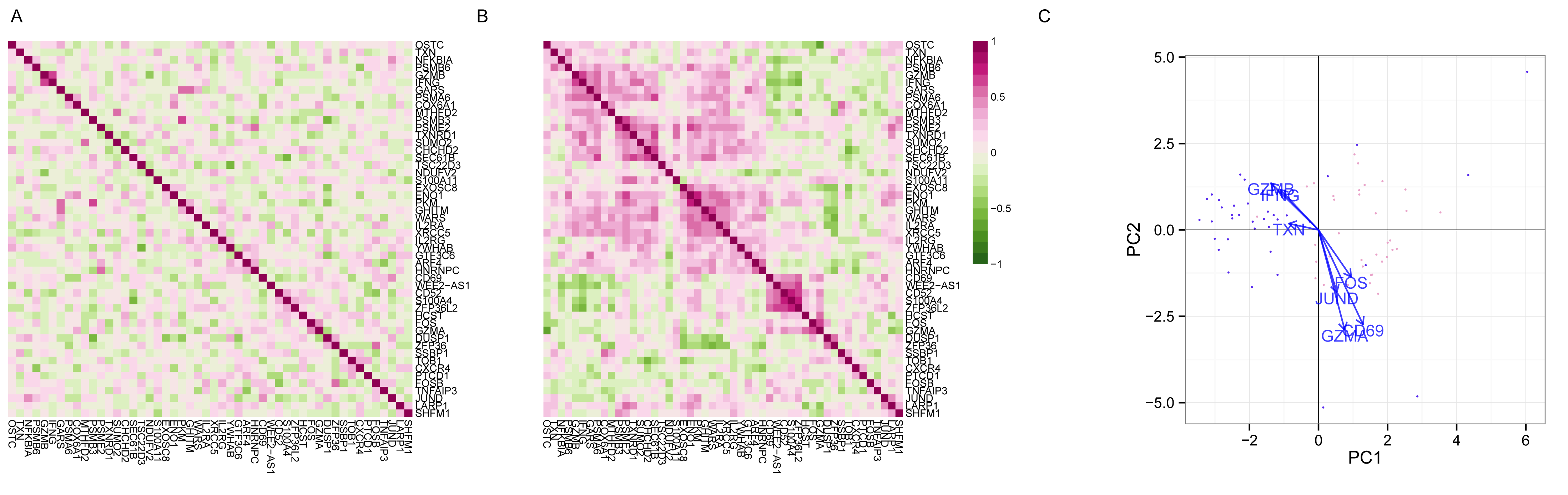
Gene-gene correlation (Pearson’s rho) of model residuals in non-stimlated (A) and stimulated (B) cells, and principal components analysis biplot of model residuals (C) on both populations using the top 50 marginally differentially expressed genes. As marginal changes in the genes attributable to stimulation and CDR have been removed, clustering of subpopulations in (C) indicates co-expression of the indicated genes on a cellular basis.

In order to assess the type-I error rate of our approach, we also applied MAST to identify differentially expressed genes across random splits of the non-stimulated MAIT cells. As expected, MAST did not detect any significant differences (Supplementary Figure 5A), whereas DEseq and edgeR, designed for bulk RNA-seq, detected large number of differentially expressed genes even at very low FDR thresholds. We examined the GO enrichment of genes detected by limma or edgeR or DESeq but not MAST and found that these sets lacked significant enrichment for modules related to the treatment of interest (Supplementary Figures 5B and 6-8). MAST’s testing framework evidently has better specificity than these approaches.

**Figure 5.**
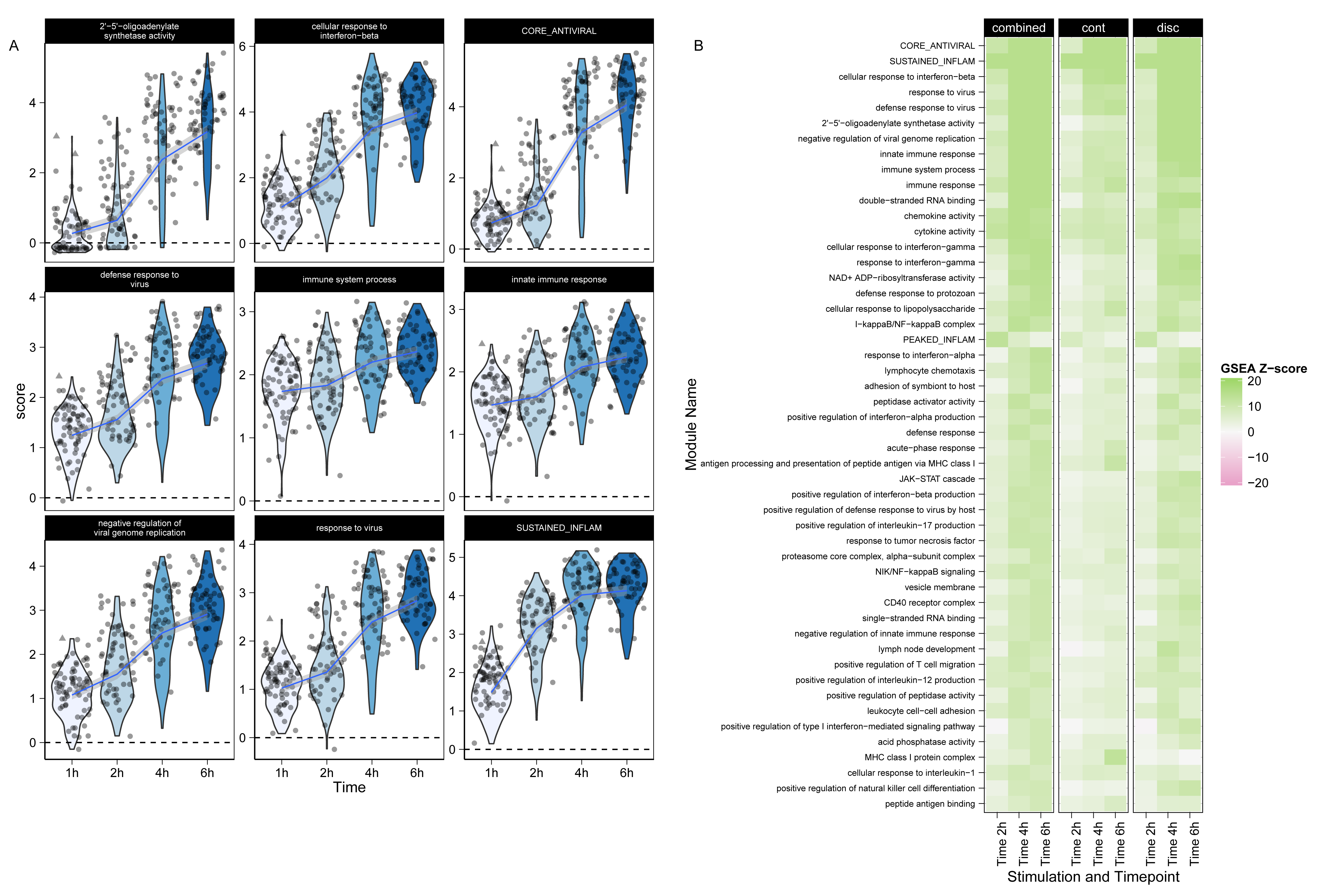
Module scores (A) and decomposed Z-scores (B) for single-cell gene set enrichment analysis for LPS stimulated cells, mDC data set, using the mouse GO biological process database. The change in single-cell module scores over time for the nine most significantly enriched modules in response to LPS stimulation are shown in A. The *core antiviral*, *peaked inflammatory* and *sustained inflammatory* modules are among the top enriched modules, consistent with the original publication. Additionally we identify GO modules *cellular response to interferon-beta* and *response to virus*, which behave analogously to the core antiviral and sustained inflammatory modules. No GO analog for the *peaked inflammatory* module was detected. The majority of modules detected exhibit enrichment relative to the 1h time point (thus increasing with time). The “early marcher” cells identified in the original publication are highlighted here with triangles. We show the top 50 most significant modules (B). The combined Z-score summarizes the changes in the discrete and continuous components of expression.

Figure 2A shows the single-cell expression (log_2_-TPM) of the top 100 genes identified as differentially expressed between cytokine (IL18, IL15, IL12) stimulated (purple) and non-stimulated (pink) MAIT cells using MAST. Following stimulation with IL12/15/18, we observe increased expression in genes with effector function including Interferon–γ (IFN– γ), granzymeB (GZMB) as has been reported in NK, NKT and memory T cells, and a concomitant downregulation of the AP-1 transcription factor network. CD69 is an early and only transient marker of activation that can be induced by stimulation of the T cell receptor or by cytokine signals. Its downregulation at the mRNA level after 24h is likely preceding subsequent protein-level downregulation^16-18^.

We used these lists of up- and down-regulated genes to define a MAIT activation score that differentiates between stimulated and non-stimulated MAITs as shown in Figure 2B. This score (see Methods), for each cell, is defined as the expected expression level across genes in a module (based on the model fit) corrected for nuisance factors (such as CDR, see Methods). The score enables us to cleanly differentiate stimulated and non-stimulated cells, and demonstrates that the stimulated MAIT population is much more heterogeneous in its expression phenotype. In particular, a few stimulated MAIT cells (SC08, SC54, SC48, SC15, SC46, and SC61 in Figure 2A) exhibit low expression of IFN–*γ* response genes, suggesting these cells did not fully activate despite stimulation. Post-sort experiments via FCM show that the sorted populations were over 99% pure MAITs (Supplementary Figure 9A), and exhibited a change in cell size upon stimulation (Supplementary Figure 9B), and that up to 26% of stimulated MAITs didn’t express IFN-γ or GZMB following cytokine stimulation (Supplementary Figure 9C). The non-responding cells in the RNA-seq experiment likely correspond to these non-responding cells from the flow cytometry experiment, and the observed frequencies of these cells in the RNA-seq and flow populations are consistent with each other (Pr(observing 6 or fewer non-responding cells) = 0.16 under binomial sampling). We discuss this heterogeneity in a further section. Importantly, the lists of up- and down-regulated genes can be used to define gene sets for gene set enrichment analysis in order to identify transcriptional changes related to MAIT activation in bulk experiments.

### Gene set enrichment analysis highlights pathways implicated in MAIT cell activation

We used MAST to perform gene set enrichment analysis (GSEA, see methods) in the MAIT data using the blood transcriptional modules of Li et al^19^. The cell-level scores for the top 9 enriched modules (Figure 3A) continue to show significant heterogeneity in the stimulated cells, particularly for modules related to T-cell signaling, protein folding, proteasome function, and the AP-1 transcription factor network. Enrichment in stimulated cells (green) and non-stimulated cells (pink) is displayed for each module for the discrete and continuous components of the model (Figure 3B, see Methods), as well as a Z-score combining the discrete and continuous parts. The enrichment in the T-cell signaling module is driven by the increased expression of IFN-γ, GZMB, IL2RA, IL2RB, and TNFRSF9, 5 of the 6 genes in the module. Stimulated cells also exhibit increased energy usage, translation and protein synthesis, while down-regulating genes involved in cell cycle growth and arrest (and other cell cycle related modules). The down-regulation of cell cycle growth inhibition genes indicates that IL-12/15/18 signals are sufficient to prepare MAIT cells for cell proliferation. Interestingly, we observe down-regulation of mRNA transcripts from genes in the AP-1 transcription factor network. This has been previously described in dendritic cells in response to LPS stimulation^20^ and, indeed, we observe this effect in the mDC data set analyzed here (Supplementary Figure 10).

Our GSEA approach is more powerful than existing methods for bulk RNA-seq data (Supplementary Figure 11), and we discover significantly enriched modules with clear patterns of stimulation-induced changes that other methods omit (Supplementary Figure 12). Two such modules include the “T-cell surface signature” and “chaperonin mediated protein folding, whose component genes show elevated expression in response to stimulation (Supplementary Figure 12A-D). These additional discoveries are not solely due to greater permissiveness in MAST. We applied MAST to identify differentially expressed gene sets across random partitions of the non-stimulated cells, to examine its false discovery rate. As expected, MAST did not detect any significant differences, which suggests that it has good type I error control.

### Residual analysis identifies networks of co-expressed genes implicated in MAIT cell activation

Much of the heterogeneity between the non-responding and responding stimulated cells remains even after removal of marginal (gene level) stimulation effects. Since, MAST models the expected expression value for each cell, we can compute residuals adjusted for known sources of variability (See Methods). The residuals can be compared across genes to characterize cellular heterogeneity and correlation. We observe co-expression in the residuals from stimulated cells that is not evident in the non-stimulated group (Figure 4A,B). Since the residuals have removed any marginal changes due to stimulation in each gene, the average residual in the two groups is comparable. The co-expression observed, meanwhile, is due to individual cells expressing these genes *dependently*, where pairs of genes appear together more often than expected under a model of independent expression.

Two clusters of co-expressed genes stand out in the residuals of the stimulated cells (Figure 4 B). These clusters show coordinated, early up-regulation of GZMB and IFN-*γ* in response to stimulation in MAIT cells and a concomitant decrease in CD69 expression, an early and transient activation marker. PCA of the model residuals highlights the non-responsive stimulated MAIT cells (Figure 4C).

Accounting for the CDR reduces the background correlation observed between genes (Supplementary Figure 13) where nearly 25% of pairwise correlations decrease after CDR correction. When the CDR is included in the model, the number of differentially expressed genes with significant correlations across cells (FDR adjusted p-value < 1%) decreases from 73 to 61 in the stimulated cells, and from 808 to 15 in non-stimulated cells. This shows that adjusting for CDR is also important for co-expression analyses as it reduces background co-expression attributable to cell volume, which otherwise results in dense, un-interpretable gene networks.

### MAST on complex experimental designs: temporal expression patterns of mouse dendritic cell maturation

Shalek et al^5^ analyzed murine bone-marrow derived dendritic cells simulated using three pathogenic components over the course of six hours and estimated the proportion of cells that expressed a gene and the expression level of expressing cells. We compared results from applying our model to those obtained by Shalek et al^5^ when analyzing their lipopolysaccharide (LPS) stimulated cells. As with the MAIT analysis, we used MAST adjusting for the CDR. MAST identified a total of 1359 differentially expressed genes (1996 omitting the CDR), and the CDR accounted for 5.2% of the model deviance in the average gene.

The most significantly elevated genes at 6h include CCL5, CD40, IL12B, and Interferon-inducible (IFIT) gene family members, while down-regulation was observed for EGR1 and EGR2, transcription factors that are known to negatively regulate dendritic cell immunogenicity^21^.

### GSEA of mouse bone marrow-derived dendritic cells

We performed GSEA with the Mouse GO modules and three modules Shalek et al^5^ identified. The blood transcriptional modules of Li et al^19^ are shown in Supplementary Figure 10. Figure 5 shows module scores for significant GSEA modules for the LPS stimulated cells where the heatmap represents Z values (see methods for details). Besides finding signatures consistent with the modules from Shalek et. al. (Figure 5A), we identify modules that show similar annotation and overlap significantly with the *core antiviral* and *sustained inflammatory* signatures, including several modules linked to type 1 interferon response and antiviral signatures (Figure 5B). The “cellular response to interferon-beta” signature (n = 22) overlaps with the original core antiviral signature (n = 99) by 13 genes (hypergeometric p = 1.24×10^−23^). The *response* and *defense response to virus* signatures overlap with the core antiviral signature by 17 of 43 and 22 of 74 genes (hypergeometric p=3.64×10^−26^ and 4.08×10^−29^, respectively), suggesting the core antiviral signature captures elements of these known signatures. The *chemokine* (n=16) and *cytokine activity* (n=51) modules overlap with the sustained inflammatory (n = 95) module by 5 and 12 genes, respectively (hypergeometric p=5.10×10^−9^ and 9.53×10^−16^). Our modeling approach identifies the two “early marcher” cells in the core antiviral module (marked with triangles on Figure 5A) corresponding to the same cells highlighted in Figure 4b of Shalek et al^5^. Other modules exhibiting significant time-dependent trends include a module of genes involved in the AP-1 transcription factor network that is down-regulated (Supplementary Figure 10), a finding which has been previously shown in human monocytes following LPS stimulation^20^. As with the MAITs, GSEA permutation analysis to evaluate type I error rates did not identify any significant modules (data not shown). These results further confirm the original findings and demonstrate the increased sensitivity of our approach. GSEA heatmaps for the other stimulations can be found in Supplementary Figure 14.

### Residual analysis of mouse bone marrow-derived dendritic cells identifies sets of co-expressed genes

We also explored stimulation-driven correlation patterns. Principal component analysis (Figure 6A) of the model residuals demonstrates a clear time trend associated with PC1, as cells increase co-expression of interferon-activated genes. After removing the marginal stimulation and adjusting for the CDR, we observe correlation between chemokines CCL5, TNF receptor CD40, and interferon-inducible (IFIT) genes (Figure 6B). A principal finding of the original publication was the identification of a subset of cells that exhibited an early temporal response to LPS stimulation. Recapitulating the original results here, when we examine the PCA of the residuals using the genes in the core antiviral module, we can identify the “early marcher” cells at the 1h time-point (Supplementary Figure 15). The co-expression plot for other stimulations can be found in the supplementary material (Supplementary Figures 16 and 17).

**Figure 6.**
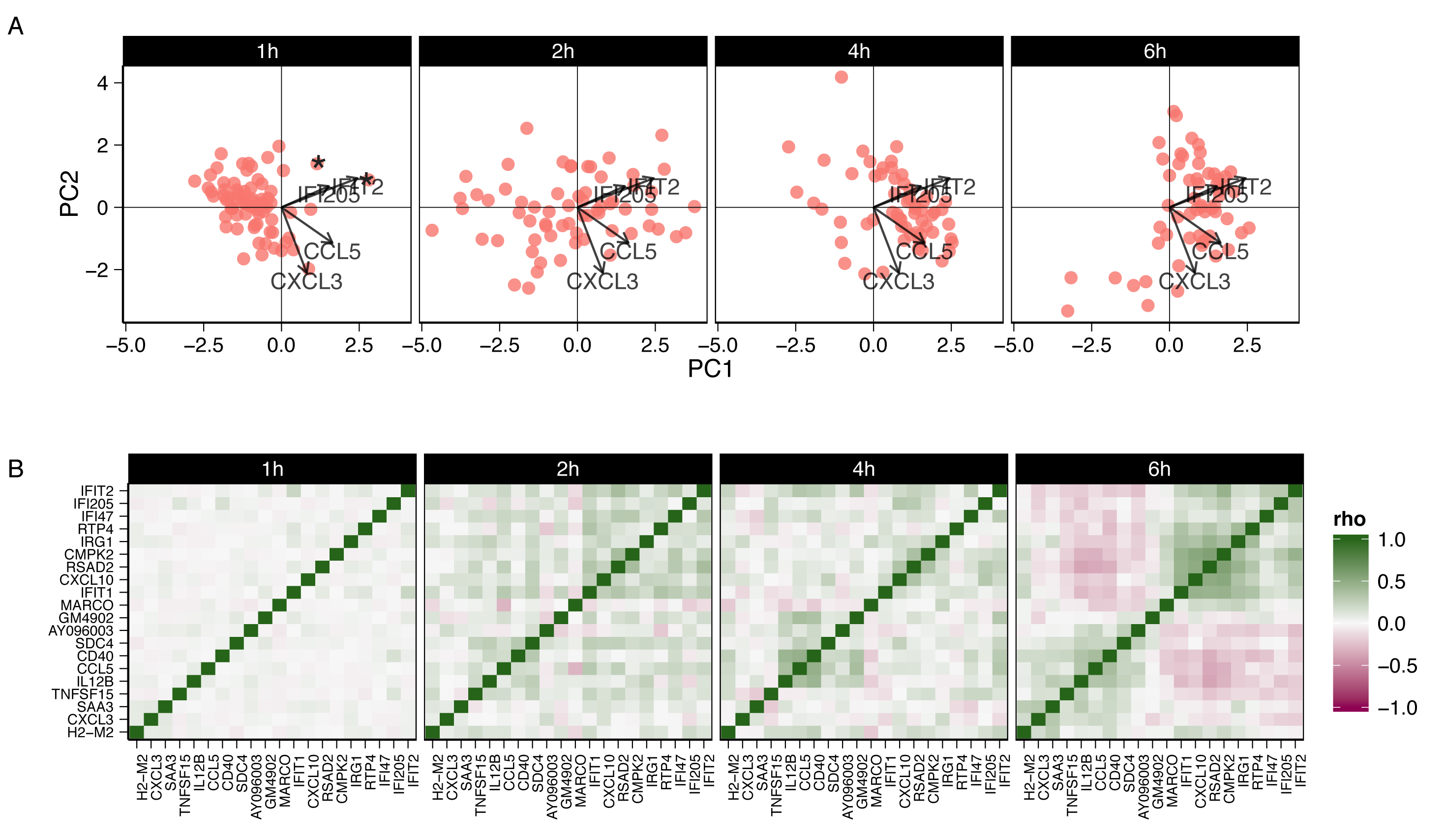
Principal components analysis biplot of model residuals (A) and Gene-gene correlation (Pearson’s R) of model residuals (B) by time point for LPS cells, mDC experiment using 20 genes with largest log-fold changes, given significant (FDR q <.01) marginal changes in expression. PC1 is correlated with change over time. The two “early marcher” cells are highlighted by an asterisk at the 1h time-point. Correlation structure in the residuals is increasingly evident over time and can be clearly observed at the 6h time-point compared to the earlier time-points.

### Discussion

We have presented MAST, a flexible statistical framework for the analysis of scRNA-seq data. MAST is suitable for supervised analyses about differential expression of genes and gene-modules, as well as unsupervised analyses of model residuals, to generate hypotheses regarding co-expression of genes. MAST accounts for the bimodality of single-cell data by jointly modeling rates of expression (discrete) and positive mean expression (continuous) values. Information from the discrete and continuous parts is combined to perform inference about changes in expression levels using gene or gene-set based statistics. Because our approach uses a generalized linear framework, it can be used to jointly estimate nuisance variation from biological and technical sources, as well as biological effects of interest. In particular, we have shown that it is important to control for the proportion of genes detected in each cell, which we refer to as the cellular detection rate (CDR), as this factor can single-handedly explain 13% of the variability in the 90% percentile gene. Adjusting for CDR at least partially controls for differences in abundance due to cell size (and other extrinsic biological and technical effects), while omitting it would lead to overestimated effects of the treatment on the system. Using several scRNA-seq datasets, we showed that our approach provides a statistically rigorous improvement to methods proposed by other groups in this context^5^.

Because our approach is regression-based, it can be used to compute residuals to explore cellular heterogeneity and gene-gene correlations after selected technical and/or biological effects have been removed. In particular, using this approach, we identify MAIT cells that do not have a typical activated expression profile in response to stimulation (Figures 2 and 3). The proportion of these cells detected in the scRNASeq experiment is consistent with what was detected in the flow cytometry experiment. These cells do not produce IFN-*γ* or GZMB upon to cytokine stimulation and exhibit expression profiles intermediate to non-stimulated and stimulated cells (Supplementary Figure 18C). The cells exhibit lower levels of IFN-*γ* and GZMB than activated cells (Supplementary Figure 18A), but also exhibit decreased expression of AP-1 component genes Fos and FosB, consistent with other stimulated cells (Supplementary Figure 18B).

As discussed by Padovan-Merhar et al^10^, care must be taken when interpreting experiments where the system shows global changes in CDR across treatment groups, as this could result in confounding treatment effect with differences in cell volume, which are not necessarily of biological interest. Our approach addresses this issue as MAST allows joint modeling of CDR and treatment effects, so the interpretation of the treatment effect is that the cell volume/CDR has been held constant. It is also possible to only use CDR as a precision variable by centering the CDR within each treatment groups, which makes the CDR measurement orthogonal to treatment. This would implicitly assume that the observed changes are treatment induced, while still modeling the heterogeneity in cell volume within each treatment group. An alternative approach would be to estimate the CDR coefficient using a set of control genes assumed to be treatment invariant, such as housekeeping or ERCC spike-ins^22,23^ and including it as an offset to the linear predictors in the regression. An analogous approach is undertaken by Buettner et. al.^22^, however it does not account for bimodality and does not jointly model technical and biological effects.

MAST is available as an R package (http://www.github.com/RGLab/MAST, doi: 10.5281/zenodo.18539). All data and results presented in this paper – including code to reproduce the results – are available at: (http://github.com/RGLab/MASTdata/archive/v1.0.0.tar.gz, doi: 10.5281/zenodo.18540). It should also be noted that while most of the methodology presented here was developed for scRNA-seq, it should be applicable to other single-cell gene expression platforms.

## METHODS

### Data Sets

Data for the MAIT study were derived from a single donor who provided written informed consent for immune response exploratory analyses. The study was approved by the relevant institutional review boards.

#### MAIT cell isolation and stimulation

Cryopreserved PBMC were thawed and stained with Aqua Live/Dead Fixable Dead Cell Stain and the following antibodies: CD3, CD8, CD4, CD161, Vα7.2, CD56 and CD16. CD8^+^ MAIT cells were sorted as live CD3^+^CD8^+^ CD4^-^CD161^hi^Vα7.2^+^ cells and purity was confirmed by post-sort FACS analysis. Sorted MAIT cells were divided into aliquots and immediately processed on a C1 Fluidigm machine or treated with a combination of IL-12 (eBioscience), IL-15 (eBioscience), and IL-18 (MBL) at 100ng/mL for 24 hours followed by C1 processing.

#### C1 processing, Sequencing, and Alignment

After flow sorting, single cells were captured on the Fluidigm™ C1 Single-Cell Auto Prep System (C1), lysed on chip and subjected to reverse transcription and cDNA amplification using the SMARTer® Ultra™ Low Input RNA Kit for C1 System (Clontech). Sequencing libraries were prepared using the Nextera XT DNA Library Preparation Kit (Illumina) according to C1 protocols (Fluidigm). Barcoded libraries were pooled and quantified using a Qubit® Fluorometer (Life Technologies). Single-read sequencing of the pooled libraries was carried out either on a HiScanSQ or a HiSeq2500 sequencer (Illumina) with 100-base reads, using TruSeq v3 Cluster and SBS kits (Illumina) with a target depth of >2.5M reads. Sequences were aligned to the UCSC Human genome assembly version 19 and gene expression levels quantified using RSEM^25^ and TPM values were loaded into R^26^ for analyses. See supplement for more details on data processing procedures.

#### Time-series stimulation of mouse bone-marrow derived dendritic cells (mDC)

Processed RNA-seq data (transcripts-per-million, TPM) were downloaded from GEO under accession number GSE41265. Alignment, pre-processing and filtering steps have been previously described^5^. Low quality cells were filtered as described in Shalek et al^5^.

### Single Cell RNA Seq Hurdle model

We model the log_2_(TPM+1) expression matrix as a two part generalized regression model. The cell expression rate given a design is modeled using logistic regression and the expression level is modeled as conditionally Gaussian given that they are expressed.

Given normalized, possibly thresholded (see supplementary material), scRNA-seq expression y = |*y*_*ig*_|, the rate of expression and the level of expression for the expressed cells are modeled conditionally independent for each gene *g*. Define the indicator Z = [*z*_*ig*_] indicating whether gene *g* is expressed in cell *i*, i.e. *z*_*ig*_ = 0 if *y*_*ig*_ = 0 and *z*_*ig*_ = 1 if *y*_*ig*_ > 0. We fit logistic regression models for the discrete variable and Gaussian linear model for the continuous variable (*Y* | *Z* = 1) independently, as follows,

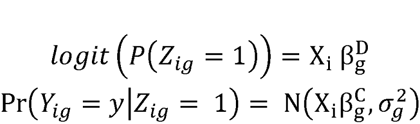

The regression coefficients of the discrete component are regularized using a Bayesian approach as implemented in the *bayesglm* function of the *arm* R package, which uses weakly informative priors^27^ to provide sensible estimates under linear separation (See supplementary material for details). We also perform regularization of the continuous model variance parameter, as described below, which helps increases robustness of gene-level differential expression analysis when a gene is only expressed in a few cells.

We define the *cellular detection rate* (CDR) as the proportion of genes detected in each cell. The CDR for cell *i* is:

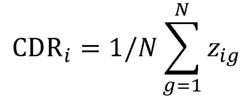

An advantage of our approach is that it is straightforward to account for CDR variability by adding the variable as a covariate in the discrete and continuous models (column of the design matrix, *X*, defined above). In the context of our hurdle model, inclusion the CDR covariate can be thought of as the discrete analog of global normalization, and as we show in the examples, this normalization yields more interpretable results and helps decrease background correlation between genes, which is desirable for detecting genuine gene co-expression.

#### Shrinkage of the continuous variance

As the number of expressed cells varies from gene to gene, so does the amount of information available to estimate the residual variance of the gene. On the other hand, many genes can be expected to have similar variances. To accommodate this feature of the assay, we shrink the gene-specific variances estimates to a global estimate of the variance using an empirical Bayes method. Let 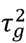 be the precision (1/variance) for *Y*_*g*_|*Z*_*g*_ = 1 in gene g. We suppose 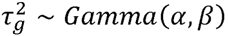, find the joint likelihood (across genes) and integrate out the gene-specific inverse variances. Then maximum likelihood is used to estimate *α* and *β*. Due to conjugacy, these parameters are interpretable providing 2*α* pseud-observations with precision *β*/*α*. This leads to a simple procedure where the shrunken gene-specific precision is a convex combination of its MLE and the common precision. This approach accounts for the fact that the number of cells expressing a gene varies from gene to gene. Genes with fewer expressed cells end up with proportionally stronger shrinkage, as the ratio of pseudo observations to actual observations is greater. Further details are available in the supplement.

### Testing for differential expression

Because *Z*_*g*_and *Y*_*g*_ are defined conditionally independent for each gene, tests with asymptotic *χ*^2^ null distributions, such as the likelihood ratio or Wald tests can be summed and remain asymptotically *χ*^2^, with the degrees of freedom of the component tests added. For the continuous part, we use the shrunken variance estimates derived through our empirical Bayes approach described above. The test results across genes can be combined and adjusted for multiplicity using the false discovery rate (FDR) adjustment^28^. In this paper, we declare a gene differentially expressed if the FDR adjusted p-value is less than 0.01 and the estimated fold-change is greater than 1.5 (on log_2_ scale).

### Gene Set Enrichment Analysis (GSEA)

Our competitive GSEA compares the average model coefficient in the *test* set (gene set of interest) to the average model coefficient in the *null* set (everything else) with a Z-test. Suppose the genes are sorted so that the first *G*_0_ genes are in the null set, and the last *G* – *G*_0_ genes are in the test set. Then, for example, to test the continuous coefficients in the gene set, the sample means of the coefficients in the test and null sets are calculated, that is, calculate 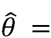 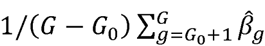 and 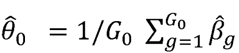. The sampling variance of 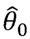, in principle, is equal to 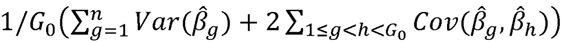, and similarly for 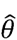.

Given this sampling variance, a Z test can be formed by comparing 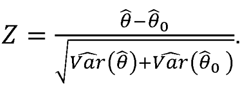

We estimate 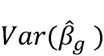 and 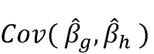 via bootstrap, to avoid relying on asymptotic approximations. In practice, we find only a few (<100) bootstrap replicates are necessary to provide stable variance-covariance estimates, however even this modest requirement can be relaxed for exploratory analysis by assuming independence across genes and using model-based (asymptotic) estimates.

Z scores are formed and calculated equivalently for the logistic regression coefficients. GSEA tests are done separately on the two components of the hurdle model and the results from the two components are combined using the Stouffer’s method^29^, which favors consensus in the two components^30^ (see supplement for details). The approach is similar to that used by CAMERA^15^ for bulk experiments in its accounting for inter-gene correlation that is known to inflate the false significance (type-I error) in permutation-based GSEA protocols^15^, although it differs in that it uses the sampling variance of each model coefficient to find the variance of the average coefficient, whereas CAMERA uses the empirical variance of the model coefficients. In our analyses we used the Emory blood transcriptional modules^19^ as well as mouse gene ontology annotations available from the Mouse Genome Informatics web site^32^.

#### GO Enrichment Analysis

Testing for enriched Gene Ontology terms based on list of genes was performed with the GOrilla online tool using the approach of comparing an unranked target list against a background list^33^.

### Residual Analysis

The hurdle model, in general, provides two residuals: one for the discrete component and one for the continuous component. Standardized deviance residuals are calculated for the discrete and continuous component separately, and then we combine the residuals by averaging them. If a cell is unexpressed, then its residual is missing and it is omitted from the average. See the supplement for details.

### Module Scores

In order to assess the degree to which each cell exhibits enrichment for each gene module, we use quantities available through our model to define module “scores”, which are defined as the observed expression corrected for CDR effect, analogous to those defined by Shalek et al^5^. The score *s*_*ij*_ for cell *i* and gene *j* is defined as the observed expression corrected for the CDR effect: *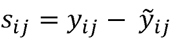 where 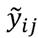* is the predicted effect from the fitted model that excludes thre treatment effects of interest. This can be interpreted as correcting the observed expression of gene *j* in cell *i* by subtracting the conditional expectation of nuisance effects. In our two part model, 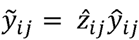 where 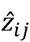 and 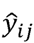 are the predicted values from the discrete and continuous components of our hurdle model.

A gene module score for cell I is the average of the scores for the genes contained in the module, i.e. ∑_{*j* ∈ *module*}_*s*_*ij*_/*|module|*

## Author Contributions

GF, AM, MY and RG developed the statistical methods, and wrote the manuscript. AM, MY, and GF wrote the R package and performed data analysis. CKS, HWM, and MP designed and performed the MAIT cell experiments and contributed to data interpretation and provided manuscript feedback. VG and PL coordinated the collection of the single cell sequencing data and contributed manuscript preparation and feedback and data interpretation. AKS contributed the mDC data and contributed manuscript feedback and to data interpretation. JD contributed to data analysis of the single-cell expression data. MJM contributed samples and to study design.

## Acknowledgements

This work was supported by NIH grants DP2 DE023321 (to M.P.) and R01 EB008400 (to R.G.). We thank all study volunteers and the Seattle HIV Vaccine Trials Unit for providing samples. We thank the James B. Pendleton Charitable Trust for their generous equipment donation. GF is an ISAC scholar.

## Reference

1. Elowitz, M. B., Levine, A. J., Siggia, E. D. & Swain, P. S. Stochastic gene expression in a single cell. Science 297, 1183–1186 (2002).

2. Raj, A., van den Bogaard, P., Rifkin, S. A., van Oudenaarden, A. & Tyagi, S. Imaging individual mRNA molecules using multiple singly labeled probes. Nature methods 5, 877–879 (2008).

3. Sanchez, A. & Golding, I. Genetic determinants and cellular constraints in noisy gene expression. Science 342, 1188–1193 (2013).

4. McDavid, A. et al. Data exploration, quality control and testing in single-cell qPCR-based gene expression experiments. Bioinformatics 29, 461–467 (2013).

5. Shalek, A. K. et al. Single-cell RNA-seq reveals dynamic paracrine control of cellular variation. Nature 510, 263–269 (2014).

6. Kharchenko, P. V., Silberstein, L. & Scadden, D. T. Bayesian approach to single-cell differential expression analysis. Nature methods 1–5 (2014). DOI: 10.1038/nmeth.2967

7. Kaufmann, B. B. & van Oudenaarden, A. Stochastic gene expression: from single molecules to the proteome. Current opinion in genetics |& development 17, 107–112 (2007).

8. Marinov, G. K. et al. From single-cell to cell-pool transcriptomes: stochasticity in gene expression and RNA splicing. Genome research 24, 496–510 (2014).

9. Marguerat, S. et al. Quantitative analysis of fission yeast transcriptomes and proteomes in proliferating and quiescent cells. Cell 151, 671–683 (2012).

10. Single Mammalian Cells Compensate for Differences in Cellular Volume and DNA Copy Number through Independent Global Transcriptional Mechanisms. 58, 339–352 (2015).

11. Voom: precision weights unlock linear model analysis tools for RNA-seq read counts. 15, R29 (2014).

12. Robinson, M. D., McCarthy, D. J. & Smyth, G. K. edgeR: a Bioconductor package for differential expression analysis of digital gene expression data. Bioinformatics 26, 139–140 (2010).

13. Anders, S. & Huber, W. Differential expression analysis for sequence count data. Genome biology 11, R106 (2010).

14. Smyth, G. K. Linear models and empirical bayes methods for assessing differential expression in microarray experiments. 3, Article3 (2004).

15. Wu, D. & Smyth, G. K. Camera: a competitive gene set test accounting for inter-gene correlation. Nucleic Acids Research 40, e133 (2012).

16. Chu, T. et al. Bystander-activated memory CD8 T cells control early pathogen load in an innate-like, NKG2D-dependent manner. Cell Rep 3, 701–708 (2013).

17. Tyznik, A. J., Verma, S., Wang, Q., Kronenberg, M. & Benedict, C. A. Distinct requirements for activation of NKT and NK cells during viral infection. Journal of immunology (Baltimore, Md.: 1950) 192, 3676–3685 (2014).

18. Smeltz, R. B. Profound enhancement of the IL-12/IL-18 pathway of IFN-gamma secretion in human CD8+ memory T cell subsets via IL-15. Journal of immunology (Baltimore, Md.: 1950) 178, 4786–4792 (2007).

19. Li, S. et al. Molecular signatures of antibody responses derived from a systems biology study of five human vaccines. Nat Immunol 15, 195–204 (2014).

20. Interferon-gamma modulates the lipopolysaccharide-induced expression of AP-1 and NF-kappa B at the mRNA and protein level in human monocytes. 24, 228–235 (1996).

21. Egr2 induced during DC development acts as an intrinsic negative regulator of DC immunogenicity. 43, 2484–2496 (2013).

22. Computational analysis of cell-to-cell heterogeneity in single-cell RNA-sequencing data reveals hidden subpopulations of cells. 33, 155–160 (2015).

23. Brennecke, P. et al. Accounting for technical noise in single-cell RNA-seq experiments. Nature methods advance on, (2013).

24. Normalization of RNA-seq data using factor analysis of control genes or samples. 32, 896–902 (2014).

25. Li, B. & Dewey, C. N. RSEM: accurate transcript quantification from RNA-Seq data with or without a reference genome. BMC Bioinformatics 12, 323 (2011).

26. Gentleman, R. C. et al. Bioconductor: open software development for computational biology and bioinformatics. Genome biology 5, R80 (2004).

27. Gelman, A., Jakulin, A., Pittau, M. G. & Su, Y.-S. A Weakly Informative Default Prior Distribution for Logistic and Other Regression Models. The annals of applied statistics 2, 1360–1383 (2008).

28. Benjamini, Y. & Hochberg, Y. Controlling the false discovery rate: a practical and powerful approach to multiple testing. Journal of the Royal Statistical Society. Series B (Methodological) 57, 289–300 (1995).

29. Annotated Bibliography of Some Papers on Combining Significances or p-values. physics.data-an, (2007).

30. Combining independent p values: extensions of the Stouffer and binomial methods. 5, 496–515 (2000).

31. Molecular signatures database (MSigDB) 3.0. 27, 1739–1740 (2011).

32. Blake, J. A. et al. The Mouse Genome Database (MGD): premier model organism resource for mammalian genomics and genetics. Nucleic Acids Research 39, D842–8 (2011).

33. Eden, E., Navon, R., Steinfeld, I., Lipson, D. & Yakhini,Z. GOrilla: a tool for discovery and visualization of enriched GO terms in ranked gene lists. BMC Bioinformatics 10, 48 (2009).

